# Transient invaders can induce shifts between alternative stable states of microbial communities

**DOI:** 10.1101/659052

**Authors:** Daniel R. Amor, Christoph Ratzke, Jeff Gore

## Abstract

Microbial dispersal often leads to the arrival of outsider organisms into ecosystems. When their arrival give rise to successful invasions, outsider species establish within the resident community, which can dramatically alter the ecosystem. Seemingly less influential, the potential impact of unsuccessful invaders that interact only transiently with the community has remained largely ignored. Here, we experimentally demonstrate that such transient invasions can perturb the stability of microbial ecosystems and induce a lasting transition to an alternative stable state, even when the invader species itself does not survive the transition. First, we develop a mechanistic understanding of how environmental changes caused by such transient invaders can drive a community shift in a simple, bistable model system. Beyond this, we show that transient invaders can also induce switches between stable states in more complex communities isolated from natural soil samples. Our results demonstrate that short-term interactions with an invader species can induce lasting shifts in community composition and function.

**One Sentence Summary:** Transient invaders can cause lasting shifts in community composition and function.

## Introduction

Microbes are extremely adept at dispersal[1], [2], their migrations allowing them to colonize the most remote environments such as young ocean crust[3], medical clinic surfaces[4], or even our space stations[5], [6]. Although this notorious flow of dispersed microbes homogenize communities around the globe[1], [2], [7], countless microbial ecosystems exhibit distinctive community stability[8], [9]. As a result, microbial ecosystems can often maintain member abundances relatively unaltered over long periods of time, show characteristic patterns of succession, or remain within their characteristic operational range in the presence of moderate disturbances, e.g. dispersal disturbances[8].

Beyond a single regime of stability, increasing empirical evidence suggests that microbial ecosystems can display alternative stable states[10]–[22]. For example, the opportunistic pathogen *Clostridium difficile* can take over the gut microbiome of susceptible hosts, leading to an unhealthy, highly persistent community state[23]. Moreover, microbial ecosystems often undergo transitions between these alternative stable states[24], [25] in response to environmental disturbance, analogous to regime shifts observed in macro-ecosystems under stress [26]–[28]. Beyond sustained external stress, short-term perturbations such as exposure to antibiotics[29] or osmotic shocks[30] can also drive lasting changes in microbial communities. Among the many different stresses that can alter microbial ecosystems, however, there is still very little understanding on how microbial dispersal affects the stability of communities.

Much like other kinds of perturbations, the arrival of invader species[31]–[33] can dramatically alter the structure of the resident community. Previous studies have characterized the effects of microbial invasions according to two main community outcomes: establishment of the invader and community resilience. In the first case, invaders survive and achieve long-term establishment in the community, potentially inducing important changes in the ecosystem such as driving some members extinct[34]. Alternatively, unsuccessful invasions occur when the resident community is resilient against the invasion. In these cases, the invader, outcompeted by the resident community, goes extinct shortly after the invasion[35]–[37]. Thus, unsuccessful invasions are typically assumed to have little impact on the resident community.

The idea that transient microbiota can impact different microbial ecosystems[38], [39] has only recently begun to receive interest. In the human gut microbiome[40], for example, manifold ingested microbes are able to survive the journey through the entire gut, remaining metabolically active and responsive to the host-associated environment[41]. Most of the time, such ingested microbes merely inhabit the gut temporarily, though in some cases, they can potentially affect both the functionality and stability of the resident community[12], [40], [42]–[44]. Nevertheless, the mechanisms allowing such transient microbes to interfere with the state of the resident community are very rarely understood[45] – specifically, whether community changes would last beyond the extinction of the invader has remained unexplored. This begs a better understanding of how transient microbiota affect microbial ecosystems in the long term.

Here, we study invasion-induced transitions in laboratory microbial communities. Using a simple bistable model system composed of two antagonistic species, we show that feedbacks between microbial growth and environmental pH can explain how unsuccessful invaders drive community shifts towards alternative community compositions. We show further that analogous shifts also occur in a multi-stable community isolated from a natural soil environment, demonstrating transient invaders can impact more complex communities. Overall, our results demonstrate that a transient invader can drive a community to an alternative stable state, even if the invader does not survive the transition.

## Results

In order to explore how invasions impact multi-stable communities, we began by using a minimal model system in which two bacterial species mutually inhibit each other. As shown in our previous work, coculturing *Corynobacterium ammoniagenes* (*Ca*) and *Lactobacillus plantarum* (*Lp*) under a serial dilution protocol leads to two alternative outcomes. *Lp* can outcompete its partner by modifying the pH towards acidic values in which *Ca* cannot grow. Alternatively, *Ca* can induce a pH change towards alkaline values that inhibits *Lp* growth. In this way, either Ca or *Lp* can grow to dominate the ecosystem, depending on their initial relative abundance[46].

In order to determine if both outcomes correspond to stable states of the system, we cocultured these two species under serial dilutions including a low migration rate, which we applied by adding fresh cells of each species into the ecosystem after every dilution cycle (Figs. 1A and S1, Methods). Despite this migration, the ecosystem still exhibited two alternative outcomes (Fig. 1B). This indicates that the two outcomes constitute alternative stable states of the ecosystem, each of them with a basin of attraction (Fig. 1B) that allows the system to resist some degree of stress or perturbation, such as moderate migration episodes.

**Fig. 1.**
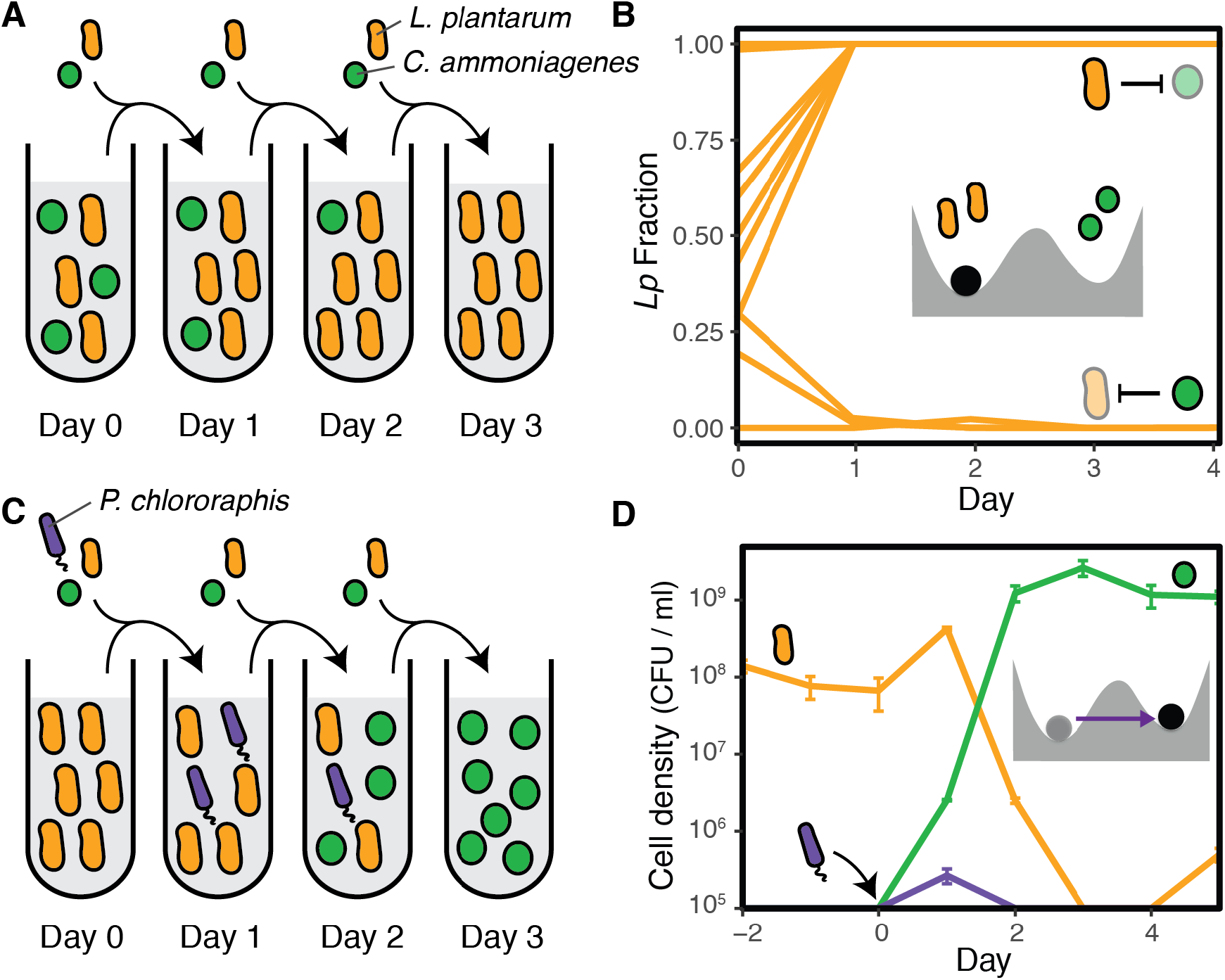
Transient invaders can induce shifts between alternative stable states in a laboratory ecosystem. **A.** We exposed cocultures of *L. plantarum* and *C. ammoniagenes* to a serial dilution protocol that includes daily migration of fresh cells from both species. **B.** Average fraction (3 replicates, standard error smaller than line width) of *Lp* cells in the community at the end of each dilution cycle, as described in A. Depending on the initial species fraction, cocultures reach a different outcome in which either species grows to dominate the system. The inset cartoon shows a mechanical analog of the ecosystem: each of the two basins of attraction can keep the marble (the community) in an alternative stable state. **C.** We explored the effects of invasions into this bistable ecosystem. The cartoon shows an unsuccessful invasion by *Pc* that nevertheless induced a shift towards an alternative stable state **D.** Time series for the cell densities during an unsuccessful invasion by *Pc* (bars show the standard error of 3 replicates). The inset cartoon depicts this invasion event as a perturbation that drives the system towards an alternative basin of stability, where it remains after the perturbation is gone.

We next studied the effects of introducing an invader species into our bistable microbial ecosystem. For this purpose, we first mixed *Lp* and *Ca* cells at a 95:5 ratio in replicate cocultures and applied 2 daily cycles of dilution and migration, allowing the system to reach one of its stable states. During the third dilution cycle, we inoculated an invader species into the coculture, simulating a single shot invasion (Fig. 1C). As expected, we observed a variety of invasion outcomes out of 7 different invader species. Only one invader candidate, namely *Bacillus cereus* (*Bc*), succeeded at establishing itself in the community (Fig. S2), while the other 6 invader species performed unsuccessful invasions (Fig. S3). *Escherichia coli* (*Ec*) or *Pseudomonas veronii* (*Pv*), for example, both mirrored classical unsuccessful invasions, as these invader species neither survived nor significantly affected the ecosystem.

Remarkably, we also identified unsuccessful invasions that induced long-term changes in the resident community, which is an invasion outcome beyond the two classical ones. In particular, an invasion by *Pseudomonas chlororaphis (Pc)* led to a shift towards the stable state governed by *Ca*. *Pc* reaches extinction (falls below the detection limit) within 48hr following its arrival into the community (Fig. 1D). Despite remaining for only a short time in the system (Fig. S4), *Pc* impacts the competition outcome between *Lp* and *Ca*, eventually leading to an increase in the *Ca* fraction that allows the community to reach its alternative stable state. This kind of unsuccessful invasion can be seen as a perturbation that drives the resident community towards the basin of attraction of an alternative stable state, where it remains after the perturbation ceases.

In addition to a change in species abundances, we observed a rapid pH shift in the microbial environment following the invader inoculation (Fig. 2A). pH measurements at the end of each daily cycle revealed consistent acidification of the media in the initial community state: the pH dropped from 6.5 to 3.7 (±0.1) during each 24hr cycle prior to the invasion. In contrast, the system persistently reached highly alkaline values (pH 9.2±0.1) in every cycle following the invader inoculation. We hypothesized that the invader could have induced the rapid shift in pH, this environmental change driving the transition between alternative states.

**Fig. 2.**
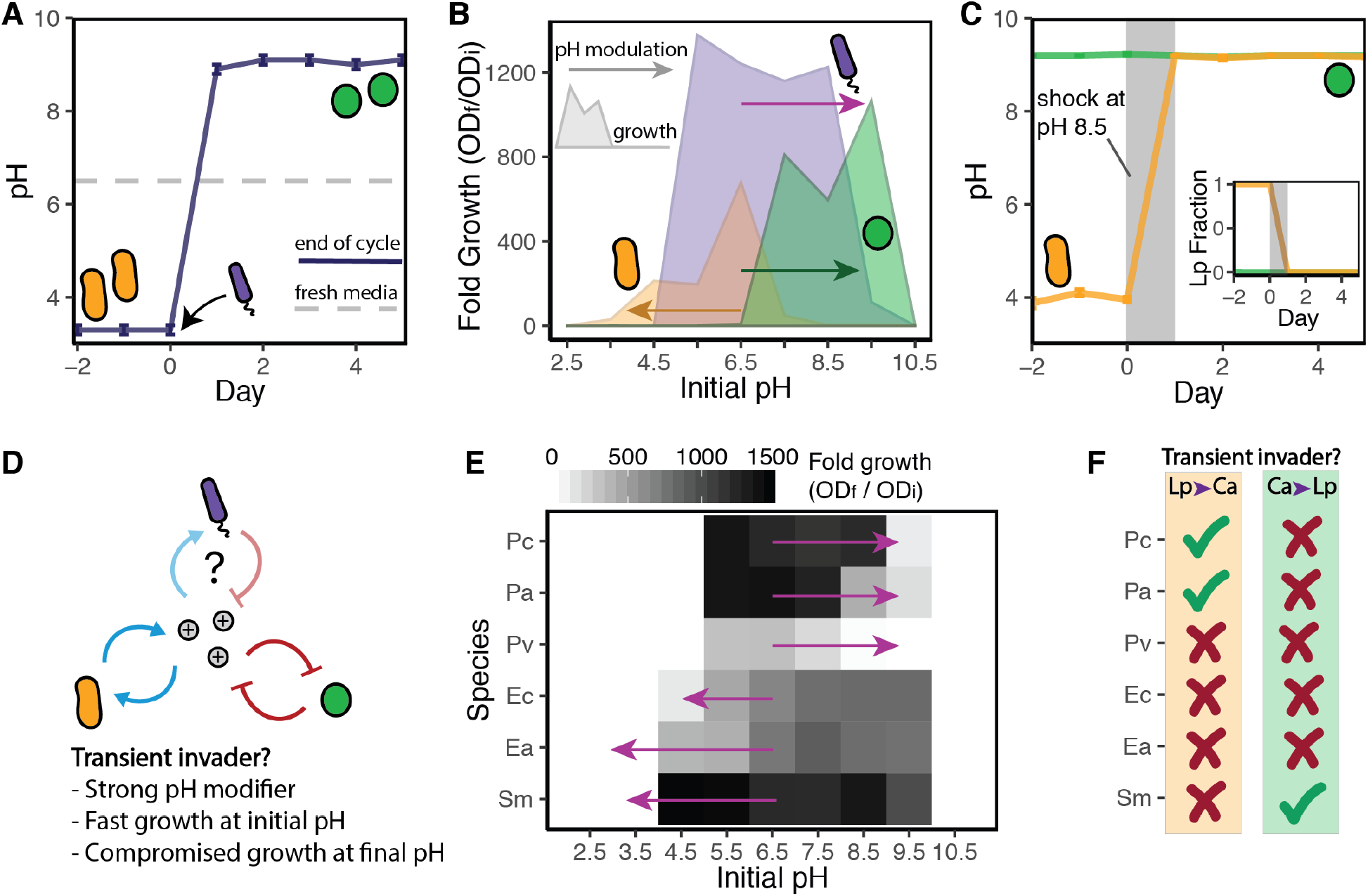
Feedback loops between microbial growth and pH can determine the community impact of transient invaders. **A.** Observed shift in pH after a *Pc* invasion into the stable state governed by *Lp*. The solid line stands for the pH at the end of each daily cycle. During each cycle, microbes induce changes in the pH of the fresh medium (dashed line) in which they were diluted into. **B.** pH range in which *Lp* (in orange), *Pc* (purple), and *Ca* (green) exhibit growth, indicated by the fold growth in OD after monocultures spent 24hr in highly (100mM phosphate) buffered media. Arrows indicate how each species modifies the pH in standard (10mM phosphate) buffer conditions. The head of the arrow points towards the pH value reached after a 24hr culture that started at pH 6.5. **C.** A temporary shock in which cells were transferred to alkaline medium during a single daily cycle (gray area) induced a transition from the *Lp* to the *Ca* state (orange), while cocultures in the *Ca* state (green) remained unaltered. **D.** Three main features observed in species that can act as transient invaders, as predicted by a minimal model that considers feedbacks between microbial growth and pH. **E.** Fold growth in highly buffered media for monocultures from 6 different species, and pH modification induced by those species in standard buffer conditions (arrows). **F.** Ticks and crosses indicate which species acted as transient invaders inducing community switches: *Pa* and *Pc* induced switches towards the alkaline state, and *Sm* was able to cause a switch towards the acidic state. Data in B and D correspond to average from 4 replicates (fold growth standard error ≤ 2, pH modification standard error ≤ 0.1).

Feedback between microbial growth and pH[46] was previously shown to drive the interaction between the resident species, *Ca* and *Lp*. Measuring the pH change after culturing *Ca* for 24hr showed that this species was able to remarkably alkalize the environment (Fig. 2B). Moreover, culturing *Ca* at different pH values revealed that its growth is strongly favored in alkaline environments, showing that *Ca* modifies the environment in a way that promotes its own growth. Similarly, *Lp* is able to acidify the environment in a way that is sustainable for itself but not for *Ca*. The invader species *Pc* displays a different pattern, since *Pc* induces a pH increase that it is not beneficial for itself (Fig. 2B). Instead, *Pc* inhibits its own growth by driving the pH towards highly alkaline values, where *Ca* grows optimally. We also studied the effects of externally applied pH shocks, which revealed that temporary perturbations in the pH were sufficient to induce transitions between the *Ca* and *Lp* stable states (Figs. 2C, S5). Together, these results show that pH modification by an invader species, namely *Pc*, can trigger switches from the stable state governed by *Lp* towards the one governed by *Ca*.

In order to better understand switches between alternative stable states that result from environmentally mediated microbial interactions, we developed a theoretical model incorporating the main features of our experimental system (Fig. S6) including daily dilutions, migration and pH modification by microbes. This minimal model uses simple step functions to determine: *i)* the pH range in which a species can actively modify the pH, and *ii)* the pH range in which each species either grows or is harmed. This simple model is able to recapitulate the observed outcomes for our bistable community, including invader-induced transitions between stable states (Fig. S6). Furthermore, we identified three specific features allowing unsuccessful invaders to cause switches between alternative stable states in *in silico* communities (Fig. 2D). In order to disrupt the initial stable state, the invader first has to be able to overcome the resident species at modifying the environment. This can be achieved through a combination of *i)* fast growth in the existing pH and *ii)* a high ability to modify the pH. As a consequence of these two requirements, the model predicts a minimum inoculum size for the invader to cause shifts in the community, a prediction that we further validated experimentally (Fig. S6). Finally, losing the competition against the resident species once the environment has changed makes the invader unsuccessful. This leads to the third requirement, *iii)* transient invaders exhibit compromised growth in the final pH. As shown in Fig. 2B, *Pc* exhibits these three generic features that enable it to act as a transient invader that alters the state of the resident community.

To determine whether these three generic features predict which invaders can trigger switches between alternative stable states, we characterized the feedback between microbial growth and pH for the 6 unsuccessful invaders mentioned previously (Fig. S3). Among those candidates, the species *Pseudomonas aurantiaca* (*Pa*) exhibited very similar features to those of the transient invader *Pc* (Fig. 2E). Indeed, an invasion by *Pa* into the *Lp* stable state also induced a community switch in the bistable ecosystem (Fig. 2F and S3). In contrast, an invasion by *Pseudomonas veronii* (*Pv*) did not result in a community switch. While *Pv* also alkalizes the environment, its lower growth rate potentially compromises its ability to overcome the initially abundant *Lp* species, as predicted by the model (Figs. 2D and S6). The last three unsuccessful invaders, namely *Enterobacter aerogenes* (*Ea*), *Escherichia coli* (*Ec*) and *Serratia marcescens* (*Sm*) modify the pH towards acidic values and, as expected based on the direction of the pH change, they were not able to induce a shift towards the alkaline (*Ca)* state. Instead, when the ecosystem was initially in the *Ca* stable state, we observed transient invasions by *Sm* that can induce a transition towards the acidic state governed by *Lp* (Fig. S7), showing that the direction of the switch can be controlled by inoculating different invader species. Analogous to the case of *Pa* and *Pch*, the transient invader *Sm* also exhibits the highest growth rates among the 3 species that induce acidification (Fig. 2E). Together, these results show that how invaders modify the environment and react to it can determine the outcome of microbial invasions in a relatively simple bistable community.

Given that we used a minimal model system that exhibits bistability between two laboratory strains, the question arises of whether analogous community dynamics could unfold from transient invasions in more complex microbial ecosystems. To address this question, we progagated a natural soil sample in the laboratory under serial dilution and looked for signatures of multistability. After 9 daily cycles, 74 out of 88 replicate cultures originating from the same soil sample exhibited optical densities above the detection limit. Analysis of the time series for the pH (Fig. 3A), as well as optical density and plating on agar suggested that the different replicates reached a wide variety of community states (Fig. S8) exhibiting different species composition.

**Fig. 3.**
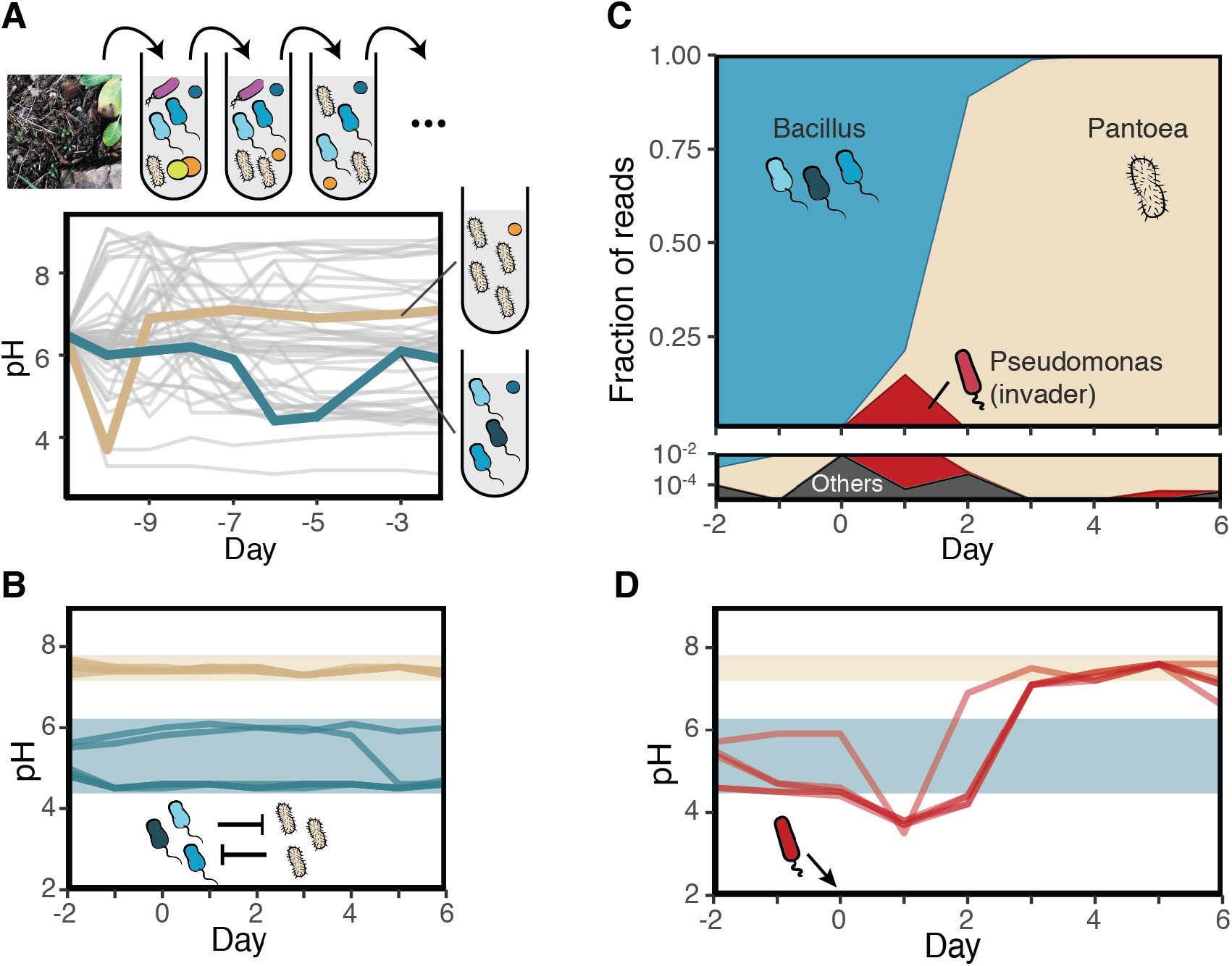
Transient invaders can drive transitions between stable states of a community isolated from the soil. Time series for the pH of 39 replicates of a soil community exposed to serial dilutions. At the end of 9 cycles, these replicates showed signs of pH stabilization (Fig. S8). The colored (blue and cream) lines correspond to two cultures in which the community composition was also stable. 16S sequencing revealed that the community in cream was highly dominated by a *Pantoea* genus, while the one in blue was governed by Bacillus (blue). **B.** Time series for the pH as the *Pantoea* and *Bacillus* communities were exposed to migration from each other. Measures for 6 technical replicates for each stable community are shown. **C.** Time series for the community composition during an invasion by *Pc* into the *Bacillus* community as revealed by 16S amplicon sequencing. The *Bacillus* community was exposed to a daily dilution protocol including migration from the *Pantoea* community as in B. **D.** Time series for the pH during the same invasion presented in **C**, along with 3 additional replicates. For reference, shaded areas indicate the observed pH range for each community in the presence of migration (as in B).

We identified two community states that were consistently able to resist low doses of migration from one another (Fig. 3B and S9). Analogous to the results in Fig. 1A and B, this indicated the presence of alternative stable states in the microcosms originated from the soil community. Sequencing revealed that one community was dominated by a *Pantoea* and the other dominated by closely related *Bacilli* strains that display a wide range of pH-growth dynamics and colony morphologies (Fig. S10).

Next, we inoculated different invader species into the two stable states governed by either *Bacillus* or *Pantoea*. Remarkably, *Pc* was consistently able to cause a switch towards the more alkaline stable state, the one dominated by *Pantoea*. Figures 3C and 3D show representative time series for the transient invasion by *Pc*, as revealed by 16S amplicon sequencing and pH measurements. Prior to the invasion, the community is dominated by the *Bacillus* genus, but after inoculation of *Pc* on Day 0 there is an increase of *Pc* accompanied by an increase in the abundance of *Pantoea.* This increase in Pantoea abundance continues even as the transient invader *Pc* is outcompeted. At the end of the experiment, *Pantoea* was dominant, *Pc* was extinct, and the abundance of *Bacillus* reads was consistent with the low migration rate applied at each dilution cycle. As before, *Pa* was also able to act as a transient invader that induces a transition to the more alkaline community state (Fig. S11). Notably, none of the considered invader species was able to induce a shift towards the stable state dominated by *Bacillus*, indicating a stronger stability of the *Pantoea* stable state against these invaders. Summarizing, these results show that transient invaders can induce shifts between alternative stable states in more complex microbial ecosystems derived from natural communities.

## Discussion

Our work demonstrates that microbial communities can experience lasting shifts in community composition induced by transient members. Analyzing both theoretically and experimentally a minimal bistable community, we have shown that feedback between microbial growth and environmental conditions (specifically, the pH) can mechanistically drive these transitions. While our results enhance the relevance of environmentally mediated interactions[46]–[49] in the microbial world, phenomenological models such as the generalized Lotka-Volterra model[50] and the stable marriage problem[51] indicate that analogous community dynamics could unfold from different microbial interactions. Analogous boom-and-bust invasions[52] have also been reported as a main cause of macroecosystem disruption. Invasive herbivores, for example, can dramatically change island landscapes before risking extinction as they deplete their main resources. Taken together, these findings suggest that transient invaders could frequently have a lasting impact also in natural microbiomes.

To date, a few longitudinal studies have linked microbial community shifts to transient invaders. Mallon *et al.* proposed that unsuccessful invaders can steer the community away from the invader’s niche[38] in a soil microbiome, although it remained unclear whether those changes lead the system to an alternative stable state. Notably, our analysis of the pH preference of the different laboratory strains is consistent with a transient invader that drives the community to an alternative niche. Although lacking strong evidence of underlying mechanisms, long-term community changes following transient invasions have also been observed in predator-prey microcosms[39], fresh water phytoplankton communities[53], as well as after infections in the human gut microbiome[44]. Our work suggests that community switches after unsuccessful invasions may not be rare, as 3 out of 6 different unsuccessful invaders induced a community switch in the minimal bistable community (2 out of 6 in the case of the soil communities). Given the currently increasing availability of temporal data sets for natural microbiomes, we expect that future analyses will frequently reveal transitions between alternative community states induced by microbial invaders.

In biomedical research, the frequent failure of probiotics[41] at establishing in the gut community has generated much skepticism about their efficacy. Our results show, however, that the manipulation of transient interactions has tremendous potential to control the long-term dynamics of microbial communities. Thus, exploring new avenues in which a set of transient microorganisms can steer the community into a healthy state[54] could lead to less intrusive interventions for the host. Beyond host-associated microbiota, future work could apply similar treatments to different microbial ecosystems (e.g., agricultural soil), and also explore the effects of co-invasions in which multiple invader species are involved.

## Acknowledgments

We thank the members of the Gore Lab for dedicated feedback. DRA specially thanks Jonathan Friedman, Gabriel Leventhal, Xiaoqian Yu, Luís Seoane and Ricard Solé for inspiring discussions. We also thank Shreyansh Umale for helping to collect the data in Fig. S10.

## Author contributions

Conceptualization: Daniel R. Amor, Jeff Gore. Data curation: Daniel R. Amor. Formal analysis: Daniel R. Amor, Christoph Ratzke, Jeff Gore. Funding acquisition: Jeff Gore. Investigation: Daniel R. Amor. Methodology: Daniel R. Amor, Christoph Ratzke. Project administration: Daniel R. Amor, Christoph Ratzke, Jeff Gore. Resources: Daniel R. Amor. Software: Daniel R. Amor. Supervision: Jeff Gore. Validation: Jeff Gore. Visualization: Daniel R. Amor. Writing – original draft: Daniel R. Amor, Christoph Ratzke, Jeff Gore.

## Data and materials availability

All data is available in the main text or the supplementary materials.

## Supplementary Materials

Materials and Methods

Figures S1-S11

References (*37-39*)

