## Supplementary material for "Transient invaders can induce shifts between alternative stable states of microbial communities": Methods and Supplementary Materials

**This PDF file includes:**

Materials and Methods
Figs. S1 to S11

### Materials and Methods

Media, Buffer, Cultures and Plating:

Overnight pre-cultures of laboratory strains were performed in Nutrient Media[1]: 10g/l of yeast extract and 10g/l of soytone (both Becton Dickinson, Franklin Lakes, USA), 10mM Sodium phosphate, pH7. The experiments were performed in Base Media[1] (hereafter, BM) supplemented with glucose and urea. The stock of base media was prepared as: 1g/l yeast extract, 1 g/l soytone, 10mM sodium phosphate buffer, 0.1 mM $CaCl_2$, 2 mM $MgCl_2$, 4 mg/l $NiSO_4$ and 50 mg/l of $MnCl_2$, 1X Trace Metals Mixture (Teknova, Hollister, CA), pH 6.5. A standard concentration of 10mM sodium phosphate in BM was used in all the experiments; with the exception of experiments measuring the growth dependence on the pH (Figure 2D) of monocultures, in which BM contained 100mM sodium phosphate. Supplemented Base Media (SBM) for the experiments was prepared daily by adding 10g/l glucose and 8g/l urea to BM. All media were filter sterilized using VWR Bottle Top Filtration Unit (VWR, Radnor, USA).

Before plating for CFU counting, experimental cultures were diluted in phosphate buffered saline (PBS, Corning, New York, USA). Plating was performed on Tryptic Soy Broth (Teknova, Hollister, USA) with 2.5% agar (Becton Dickinson, Franklin Lakes, USA) in which we adjusted the pH to different values for selective plating (see below).

Overnight pre-cultures took place in 5ml NM, inside 50mL Falcon Tubes for 24hr in the case of Ca and Lp, and 16hr for the rest of strains, shaking at 250 rpm on a New Brunswick Innova 2100 shaker (Eppendorf, Hauppauge, NY). Experimental cultures took place in 96-deepwell plates covered with AearaSeal adhesive sealing films (Excell Scientific, Victorville, USA), shaking at 1350 rpm pm a Heidolph platform shaker (Titramax 100, Heidolph North America, Elk Grove Village, IL). All pre-cultures and cultures were incubated at 30C, relative humidity 50%.

Laboratory strains

*Lactobacillus plantarum* (ATCC 8014), *Corynebacterium ammoniagenes* (ATCC 6871), *Pseudomonas chlororaphis* (*ATTC 9446* ), *Enterobacter aerogenes* (ATCC 13048), *Pseudomonas aurantiaca* (ATCC 33663), *Pseudomonas veronii* (ATCC 423 700474), *Serratia marcescens* (ATCC 13880) were obtained from ATCC. *Bacillus cereus* was obtained from Ward's Scientific Catalog. *Escherichia coli* MC4100 (CGSC #6152) was obtained from the E. coli Genetic Stock Center.

Soil microcosms

The soil was sampled from a lawn in Cambridge Massachusetts, at depth ~15 cm. About 20 grains of soil (~0.5g) were diluted into 20 ml of PBS, vortexed at intermediate speed for 30s and then incubated on shaker at 250rpm. After 30 minutes, the sample was allowed to settle for 5 minutes and the supernatant was transferred to a new Falcon tube. 7μL aliquots of this supernatant were then transferred into 203 μL of SBM in a 96-deepwell plate, inoculating a total of 88 replicates. We applied 7 daily dilution cycles to the 88 replicates as explained in the main text, and at the end of day 7 (which corresponds to day -4 in Figure 3A) we froze the resulting communities (transferring them to a final concentration of 40% glycerol, then storing at -80C).

Invasions to the soil communities were initiated by sampling the -80C stocks to inoculate 203μL of SBM in 96-deepwell plates. Then we exposed these communities to daily dilution cycles with migration (see Migration below). To facilitate the thawed samples to recover stability after coming from -80C, migration was not applied during the first daily cycle. Invaders were inoculated along the dilution protocol after the end of the 4$^{th}$ cycle. We used *Ea, Ec, Pa, Pc, Pv,* and *Sm* as candidate invader species for the soil communities.

Daily dilutions

At the end of every daily cycle, a 30 fold dilution was applied by transferring 7μL of the experimental cultures into 203μL of fresh SBM using a Viaflo 96-well pipettor (Viaflo settings: Pipette-Mix program, aspirating 7μL, 3 mixing cycles, mixing volume 10μL, speed 6).

Migration

Overnight cultures of both *Ca* and *Lp* were washed in 15ml of BM. To increase accuracy at adjusting the population densities of the migrant cells, we first adjusted both monocultures to an OD/cm of approximately 2.0, then performed a second round of adjustment to a final OD/cm of 0.37 for Ca and 0.24 for Lp. We then mixed 10ml of each monoculture, resulting in 20ml of the migrant cells mix. Using a Viaflo 96-well pipettor, 3μL of a fresh migrant mix was inoculated in the experimental cultures right after each dilution cycle (Viaflo settings: Pipette-Mix program, aspirating 3μL, 3 mixing cycles, mixing volume 10μL, speed 6). This resulted in a daily inoculation of $(1.2\pm0.1)\cdot10^5$ fresh cells from each species into the culture (Fig. S1).

In order to apply migrations between the stable states of the soil communities, at the beginning of the experiment we initiated several control cultures from both stable states. These control cultures were exposed to 30 fold daily dilutions in parallel to the rest of experimental cultures. During each dilution cycle, samples of these control cultures were diluted by a 100 fold in BM using the Viaflo. The resulting dilutions from each stable state were mixed at a 1:1 ratio, so that the composition of migrant cells include species from both stable states. Migration was applied by transferring 3μL of the resulting mix into wells with fresh media where the experimental cultures would be also propagated through the regular 30 fold dilution. This resulted in $4\cdot10^{-3}$ volume ratio of migrant cultures relative to the regularly diluted community at the beginning of each daily cycle.

Invader inoculation

Overnight monocultures of the strains used as invaders were washed in 15ml of BM. Then we adjusted the OD/cm to obtain a final population density of approximately $3.3\cdot10^8$ cells/mL (OD vs CFU assays were performed previously in order to guide this adjustment). During the dilution cycle indicated in each figure caption, we inoculated 3μL of the invader monocultures, resulting in an inoculation of approximately $10^5$ invader cells (<10% of the total number of cells transferred at the beginning of the cycle, Fig. S1). The same procedure was used for the experiments in Fig. S6F, followed by a series of additional 10-fold dilutions in order to obtain the reported inoculum sizes. A higher inoculum size of $3\cdot10^6$ was used in Fig. S7 for *Sm* to induce transitions towards the *Lp* state.

Characterization of the soil *Bacilli* strains

To isolate the 3 bacillus strains used in Fig. S10, we plated a diluted sample of the community (corresponding to Day -4 in Figure 3A, cream colored curve) into a TSB agar plate. We picked each of the 3 colony morphologies and performed an overnight culture in NM, then we store the monocultures (-80C, 40% glycerol). Plating samples of each overnight monoculture revealed a unique colony morphology for each monoculture, confirming a heritable colony morphology that is distinctive of each strain (Fig. S10).

For the competition experiments between the isolated Bacillus strains (Fig. S10), we grew colonies from the frozen isolates (1 day on TSB agar plates, 30C). We then picked individual colonies to perform overnight cultures in NM. The next morning we washed the monocultures in 15ml of BM and then measured their OD and adjusted to a final population density of $3.3 \cdot 10^8$ cells/mL CFU/ml (a previous, independent CFU vs OD assay was performed in order to guide the population density adjustments). Then we mixed the monocultures at the indicated ratios and inoculate 3uL of the mixes into 207uL of SBM to initiate the daily dilution cycles.

<u>Estimation of population densities (CFU/ml)</u>

For Colony Forming Units (CFU) counting, 10μL droplets of PBS-diluted cultures were plated on agar. We used agar plates at pH 5 for selective plating of Lp, pH 10 to select for Ca colonies, and pH 7 to count the abundance of the rest of the strains (including those in the soil communities). Colonies from the different invaders generally grew faster than Ca and Lp on pH7 agar, which allowed measurements of invader cell densities at very low abundances (down to relative fractions of approximately $10^{-4}$).

To prepare the 10μL droplets, we serially diluted the experimental cultures via 10 fold dilutions (maximal dilution factor was $10^{-7}$) using a 96-well pipettor (Viaflo 96, Integra Biosciences, Hudson, USA) using the program *"pipet/mix"* (pipetting volume: 20μl, mixing volume: 180μl, mixing cycles: 5, mixing and pipetting speed: 8). 10μl droplets were then transferred to 150-mm diameter agar plates with the 96-well pipettor (program *"reverse pipette"*, uptake volume: 20μl, released volume: 10μl, pipetting speed: 2). Droplets were allowed to dry and the plates were incubated at room temperature for one to two days until colonies were visible. The different dilution steps allowed to find a dilution at which colonies could be optimally counted with a Leica dissecting microscope (between ~5 and ~50 colonies). Up to 3 plating replicates per condition were performed to increase accuracy at measuring population densities.

<u>pH measurement</u>

To measure the pH of the microbial cultures, 150μl samples were transferred into 96-well PCR plates (VWR, Radnor, USA) and the pH was measured using a pH microelectrode (Orion, PerpHecT, ROSS).

<u>16S rRNA gene sequencing and analysis</u>

The DNA extractions were performed using Agencourt DNAdvance A48705 extraction kit (Beckman Coulter, Indianapolis, IN, USA) following the provided protocol. The obtained DNA was used for 16S amplicon sequencing targeting the V4-V5 region. The sequencing was done on an Illumina, MySeq by CGEB - Integrated Microbiome Resource at the Dalhousie University, Halifax, NS, Canada.

We used the R package DADA2[2] to obtain the Amplicon Sequence Variants (ASV) as described Callahan *et al*[3]. Taxonomic identities were assigned to the ASVs by using the GreenGenes Database Consortium (Version 13.8)[4] as reference database.

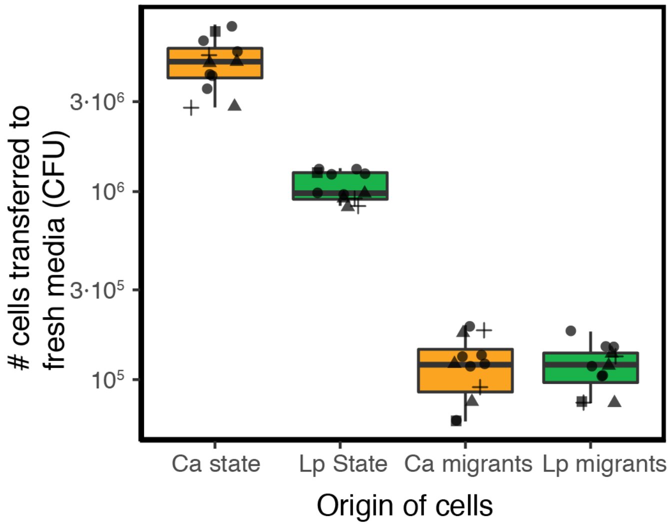


**Fig. S1.**
**Number of cells transferred to fresh media during daily dilution and migration steps.** Box plot for the number of cells transferred from the *Ca* stable state, *Lp* stable state, or inoculated as *Ca* migrants and *Lp* migrants, respectively from left to right. Different symbol shapes stand for data from independent experiments, data points with the same shape correspond to technical replicates. The two data sets on the left also indicate that the *Ca* and the *Lp* stable states had different saturation densities (which correspond to 30x the average number of cells indicated in this panel), in agreement with the different saturation densities observed in the first and last data points in Figs. 1D, S3E, S3F and S4. The applied migration rate introduces $(1.2\pm0.1)\cdot10^5$ cells/day of each species. For this migration rate, *Ca* cells migrating into the *Lp* stable state constitute up to 10% of the total number of cells transferred at the beginning of each cycle, and up to 1% for the case of *Lp* cells migrating into the *Ca* stable state.

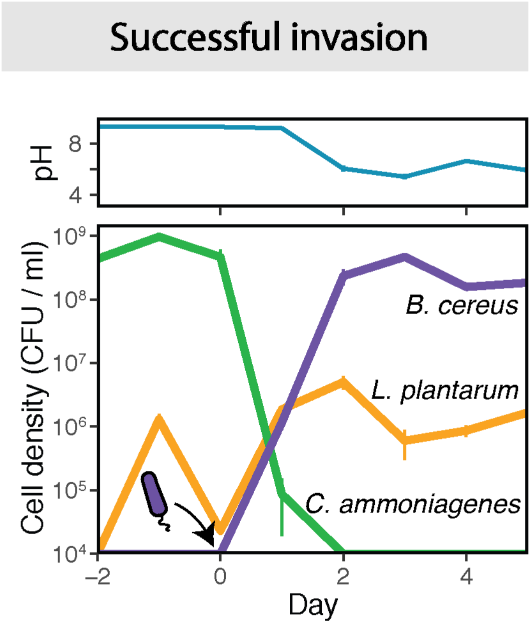


**Fig. S2.**
**Successful invasion by Bacillus cereus.** Applying daily dilution cycles including migration (Main Text), we initiated cocultures at a 5:95 ratio of *Lp* to *Ca* cells, allowing the first 3 cycles for the system to reach stability. After its inoculation at day 0, *Bc* experienced growth and was able to establish in the community while displacing *Ca*, illustrating the classical scenario of a successful invasion. The top panel shows the time series of the pH measured at the end of each cycle. Vertical bars correspond to standard error of 3 technical replicates (often smaller than the curve width).

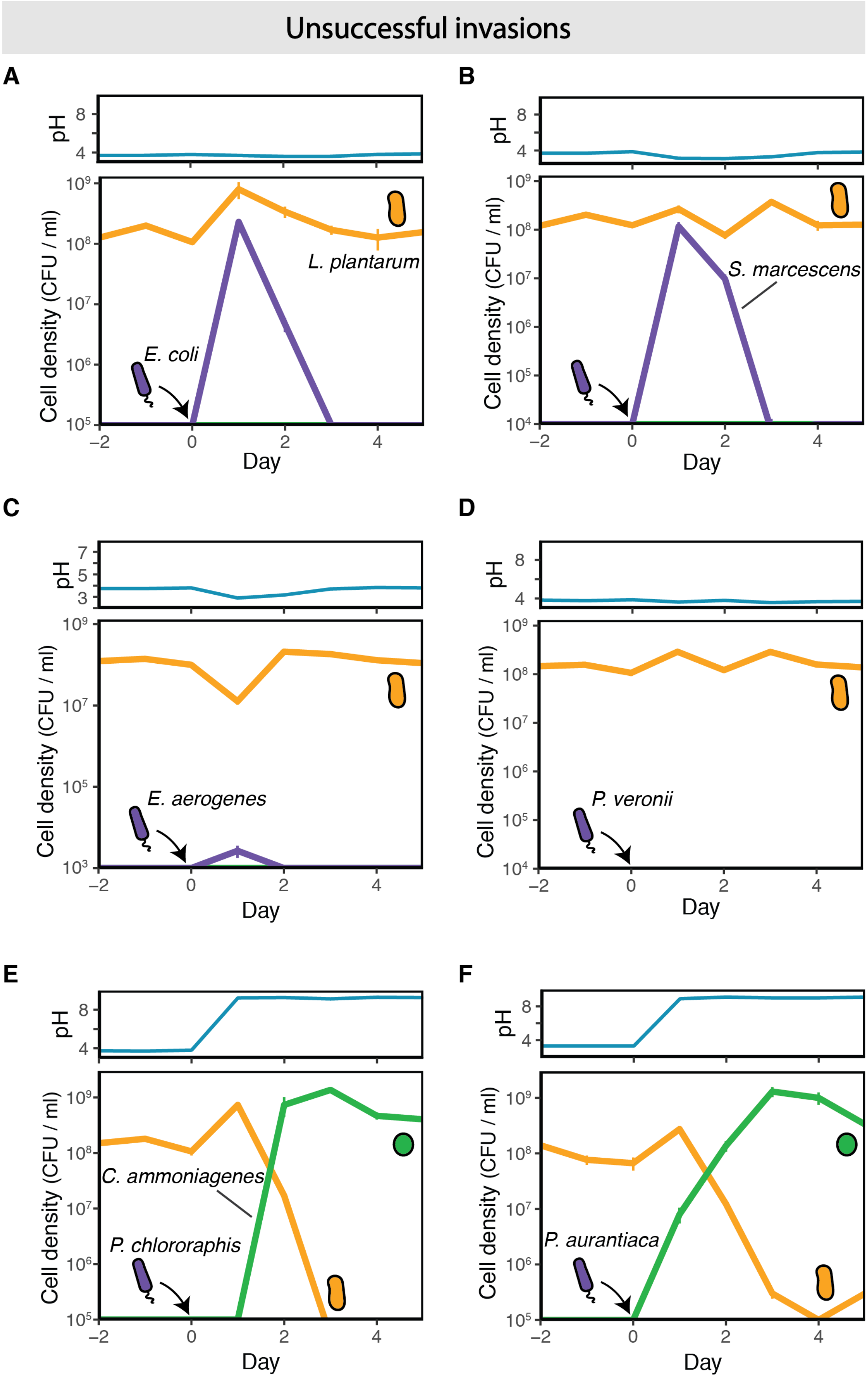

Unsuccessful invasions
A
B
C
D
E
F
pH
Cell density (CFU / ml)
Day
L. plantarum
E. coli
S. marcescens
E. aerogenes
P. veronii
C. ammoniagenes
P. chlororaphis
P. aurantiaca

**Fig. S3.**

**6 different species performed unsuccessful invasions into the *Ca* and *Lp* ecosystem.** We initiated replicate cocultures at a 95:5 ratio of *Lp* to *Ca* cells, allowing 3 initial cycles for the system to reach stability. Selective plating on agar at the end of these initial cycles revealed no CFUs from *Ca*, indicating that *Ca* cells entering via migration were not able to survive an entire 24hr cycle in the stable state governed by *Lp*. **A**. After its inoculation at Day 0, *Ec* cells experienced growth and survived in the system during two cycles. However, *Ec* experienced extinction at day 3, indicating that this species had lost the competition with the resident community, which remained in the *Lp* stable state. **B-D.** Unsuccessful invasions by *Serratia marcescens (Sm)*, Enterobacter aerogenes (*Ea),* and *Pv* species. Analogous to the case in A, these invasions mirrored classical unsuccessful invasions: the invader species experienced extinction without inducing significant change in the resident community. In D, *Pv* went extinct during the first 24hr cycle, as no CFU from this species were detected from agar plating at the end of the cycle. **E-F**. Unsuccessful invasions by *Pc* and *Pseudomonas aurantiaca (Pa)*, respectively, triggered switches from the *Lp* to the *Ca* stable state, but the invader species did not survive the transition. In all cases, time series for the cell population densities (main panels) and pH (top panels), both measured at the end of each cycle, are shown. Error bars (often smaller than the line width) in all panels show the standard error for 3 technical replicates. Data obtained from an independent experiment to the one in Fig. 1D.

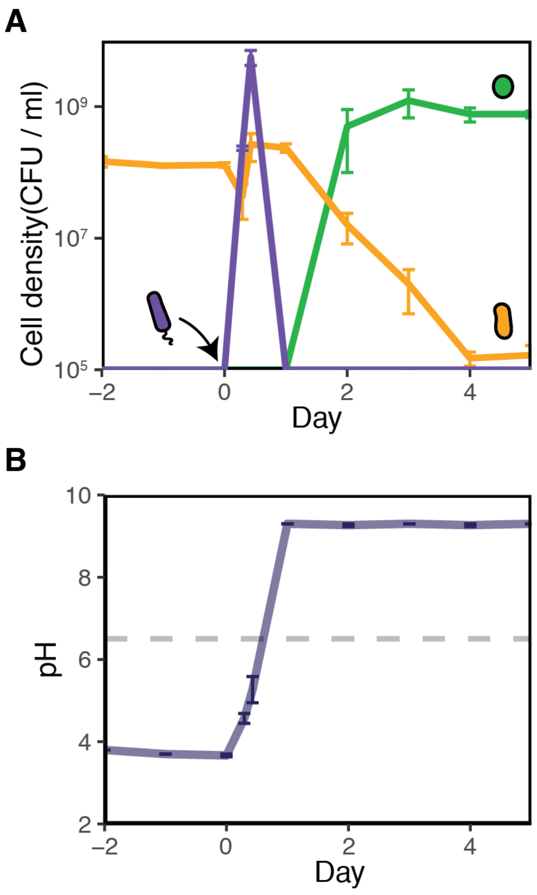


**Fig. S4.**
**The invader *Pc* thrives and decays within the 24hr following inoculation into the Lp stable state A.** Time series for the cell density of *Lp* (orange), *Ca*(green) and *Pc* (purple) during a transient invasion that induced a switch between the two alternative states. Within the first 24hr after its inoculation, *Pc* grew to high density and experienced a following decay towards extinction. **B.** Time series for the observed pH revealing a rapid shift towards alkaline values following the inoculation of *Pc*. In order to avoid any potential contamination arising from the pH measurements at 7 and 10hr post-inoculation of *Pc*, we used additional replicate cocultures that were discarded after the measurements. Error bars indicate the standard error for 3 technical replicates from an independent experiment to those in Figs. 1D and S3.

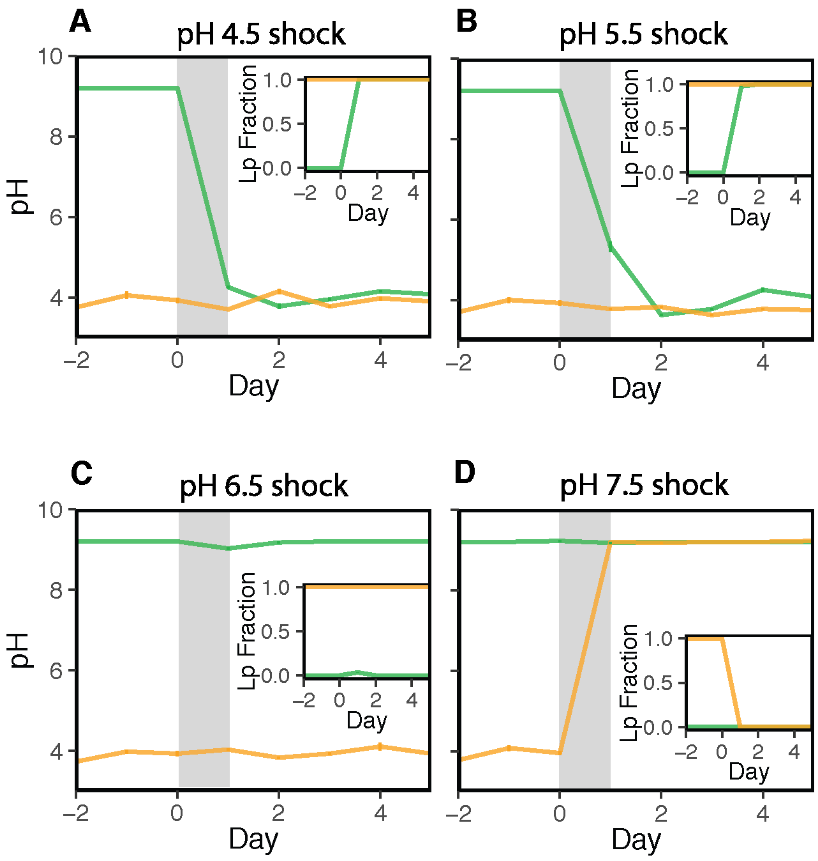


**Fig. S5.**

**Temporary perturbations in pH induce shifts between the *Lp* and the *Ca* states. A.** Starting at either 95:5 or 5:95 ratio of *Lp* to *Ca* cells, we allowed 3 cycles of dilution and migration for the system to reach the *Lp* stable state (orange) or the *Ca* stable state (green), respectively. At day 0, the cocultures were diluted into Supplemented Base Media buffered at pH 4.5 (100uM phosphate buffer) for only one cycle (shadowed area). This temporary shock induced a switch towards the *Lp* stable state in those cocultures that had previously reached the *Ca* stable state (green), while it did not significantly affect the cocultures that were already in the *Lp* state. **B.** Analogous to the previous case, a temporary shock at pH 5.5 triggered a switch from the *Ca* to the *Lp* stable state. **C.** A temporary shock at pH 6.5 did not significantly affect cocultures in either of the two stable states. **D.** A temporary shocks at pH 7.5 induced a switch from the *Lp* to the *Ca* stable state. In A-D, the main panel shows the time series for the pH measured at the end of each cycle, while the inset stands for the fraction of *Lp* cells in the system. Error bars (generally thinner than the line width) correspond to the standard error of 4 technical replicates.

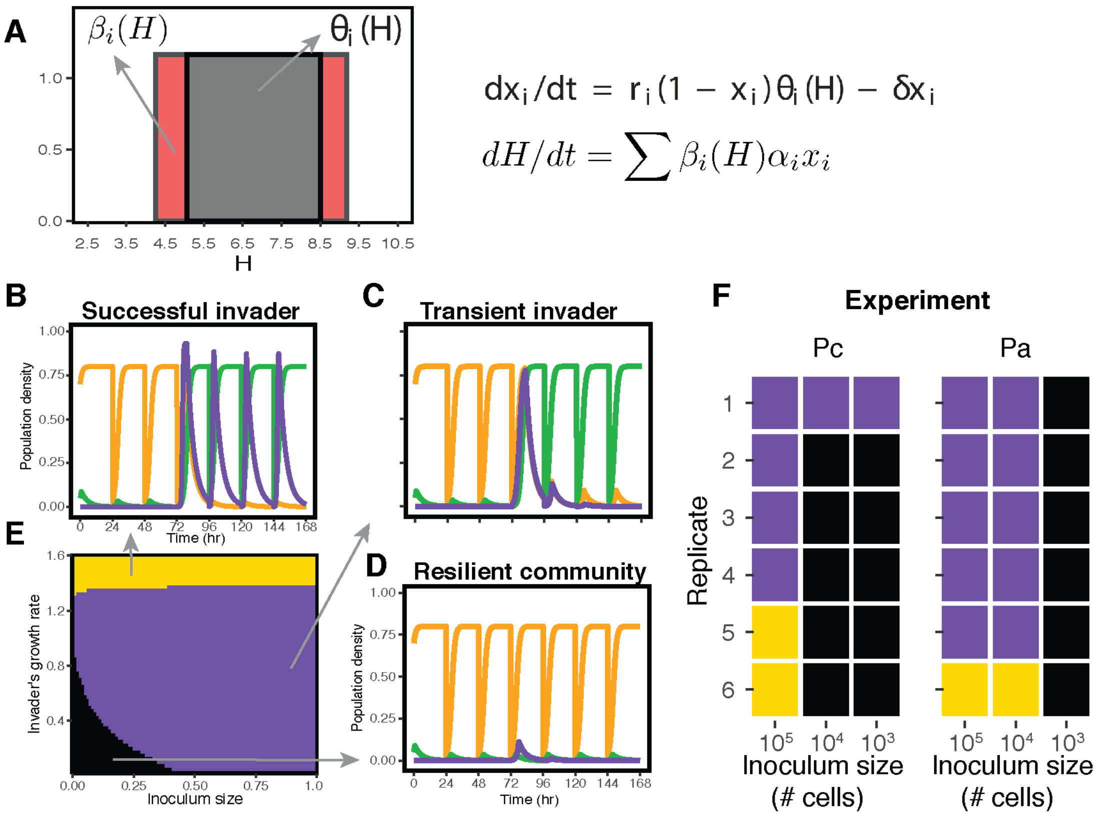


**Fig. S6.**

**Minimal model incorporating feedbacks between microbial growth and pH predicts a minimum inoculum size for the invasions to induce community shifts. A.** The minimal model considers step functions that determine the range of pH tolerance (gray area) for each species, and the range in which microbes can modify the pH (red area). The density $x_i$ of species $i$ grows logistically (with maximum growth rate $r_i$) as long as the pH lies within its tolerance range $\theta_i$. In addition, $\beta_i$ determines the range in which each species can modify the pH at a rate $\alpha_i$, as indicated by the set of equations. **B-D.** Time series for the species densities in three different scenarios, respectively: successful invasion in which the invader establishes, community shift induced by a transient invader, and resilience of the community against an unsuccessful invasion. **E.** The phase space for the three outcomes predicts a minimum inoculum size for transient invaders to induce a community shift, which lies at the interface between resilience (black region) and the shift between alternative stable states (purple). Parameter values for species 1: $r_1 = 1$, $\alpha_1 = -1$, $\theta_1 = [3.5,7.5]$, $\beta_1 = [3.8, 7.5]$. Parameter values for species 2: $r_2 = 1$, $\alpha_2 = 1$, $\theta_2 = [6,10]$, $\beta_2 = [6, 9.3]$. Parameter values for the invader species: $\alpha_3 = 1.1$, $\theta_3 = [5,8]$, $\beta_3 = [5, 8.5]$. **F.** Inoculating different initial amounts of invader cells showed that *Pc* and *Pa* require a minimum inoculum size to induce community shifts. In the experiment for the data in F, we observed some cases in which the invader had not reached extinction at the end of the experiment (2 out of 6 replicates when we inoculated $10^5$ cells of *Pc*, 1 out of 6 replicates for two different inoculum sizes of *Pa* cells).

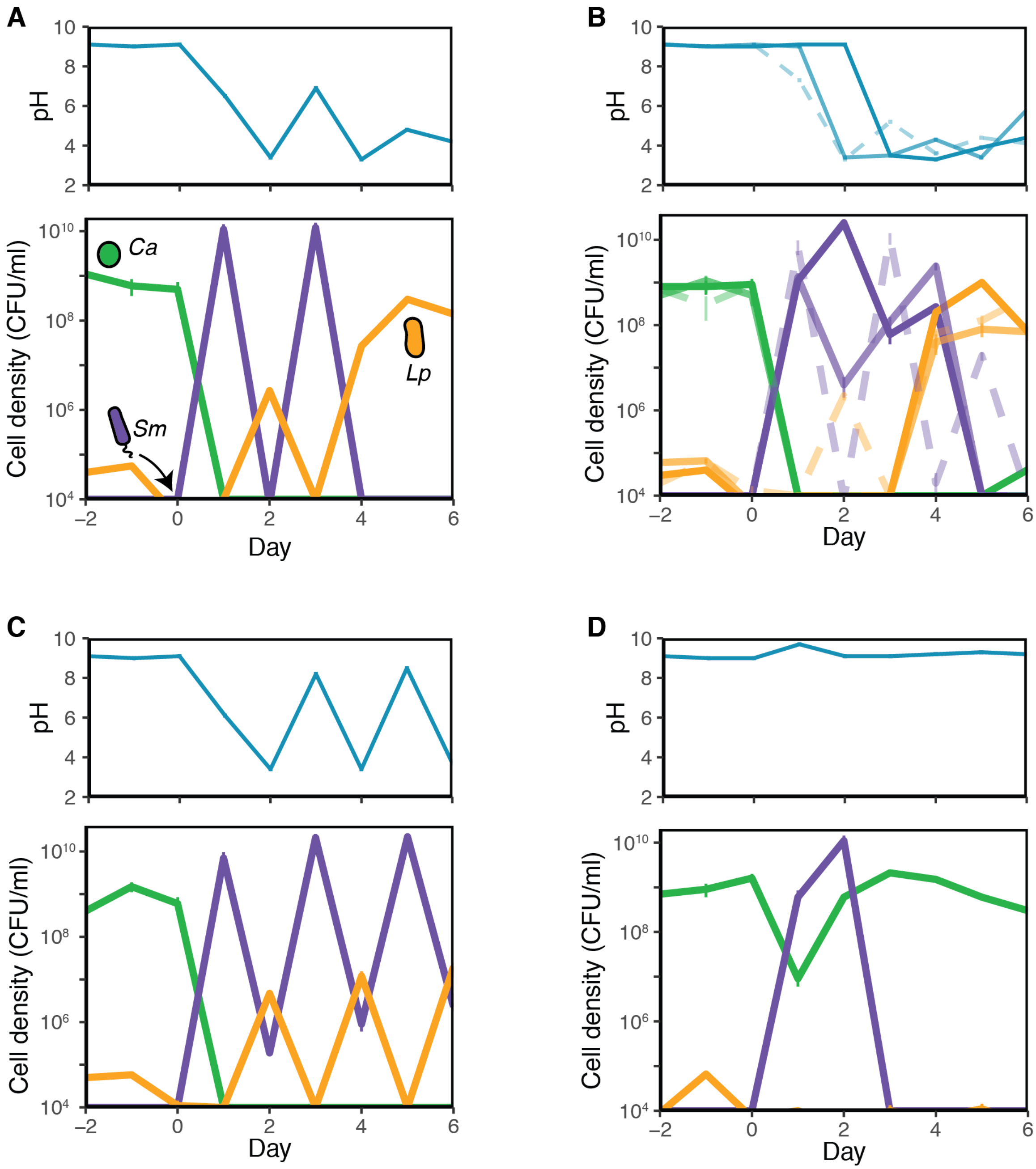


**Fig. S7.**
**The invader *Serratia marcescens* can induce transitions from the alkaline (*Ca*) to the acidic (*Lp*) stable state.** We observed a transition from the Ca to the Lp stable state in 4 out of 6 technical replicates, after inoculating *Sm* cells. **A.** Sm remained in the system for 3 cycles after its inoculation. The system reached the Lp stable state after *Sm* went extinct at Day 4. **B.** In these three replicates (solid, shade, and dashed lines) *Sm* was present in the system until Day 4 (5 in the case shown by the dashed line). No *Sm* cells were detected at the end of the experiment, and the system had transitioned into the *Lp* state. **C.** In one replicate *Sm* performed a successful invasion, as it coexisted with *Lp* until the end of the experiment. **D.** In this replicate *Sm* performed a (classical) unsuccessful invasion that did not induce a switch between stable states.

In all the main panels, the error bars correspond to the standard deviation for a Poisson distribution centered at the observed CFU for each species in a single replicate. The upper panels correspond to the pH measurements for the cultures shown in the main panels.

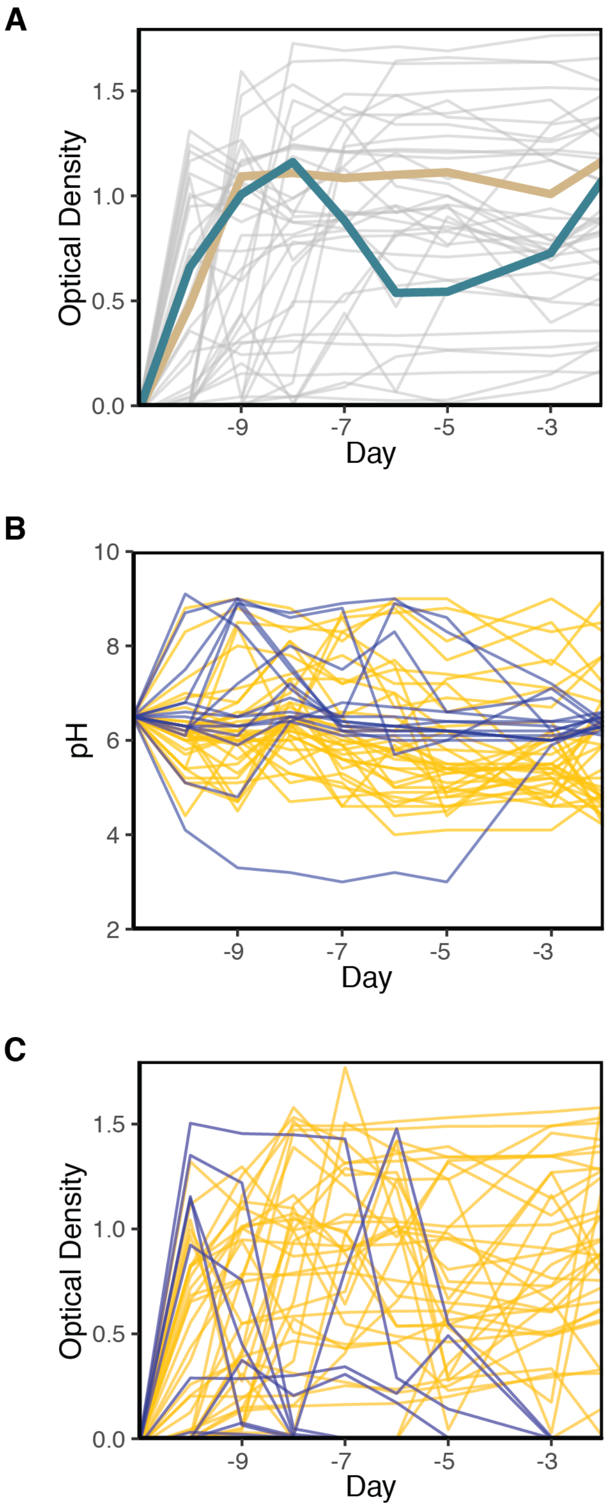


**Fig. S8.**

**Early dynamics of soil communities in laboratory environments. A.** Time series for the optical density of the soil communities. The data corresponds to the 39 cultures shown in Fig. 3A, which exhibited signatures of pH stabilization (specifically, pH change between the last two cycles <0.5). **B.** pH time series for the additional 49 cultures that were inoculated with soil samples. Communities that went extinct during the experiment are shown in blue, and communities for which the pH was not stable (pH change between the last two cycles ≥0.5) are shown in yellow. **C.** Time series for the OD of the cultures shown in B.

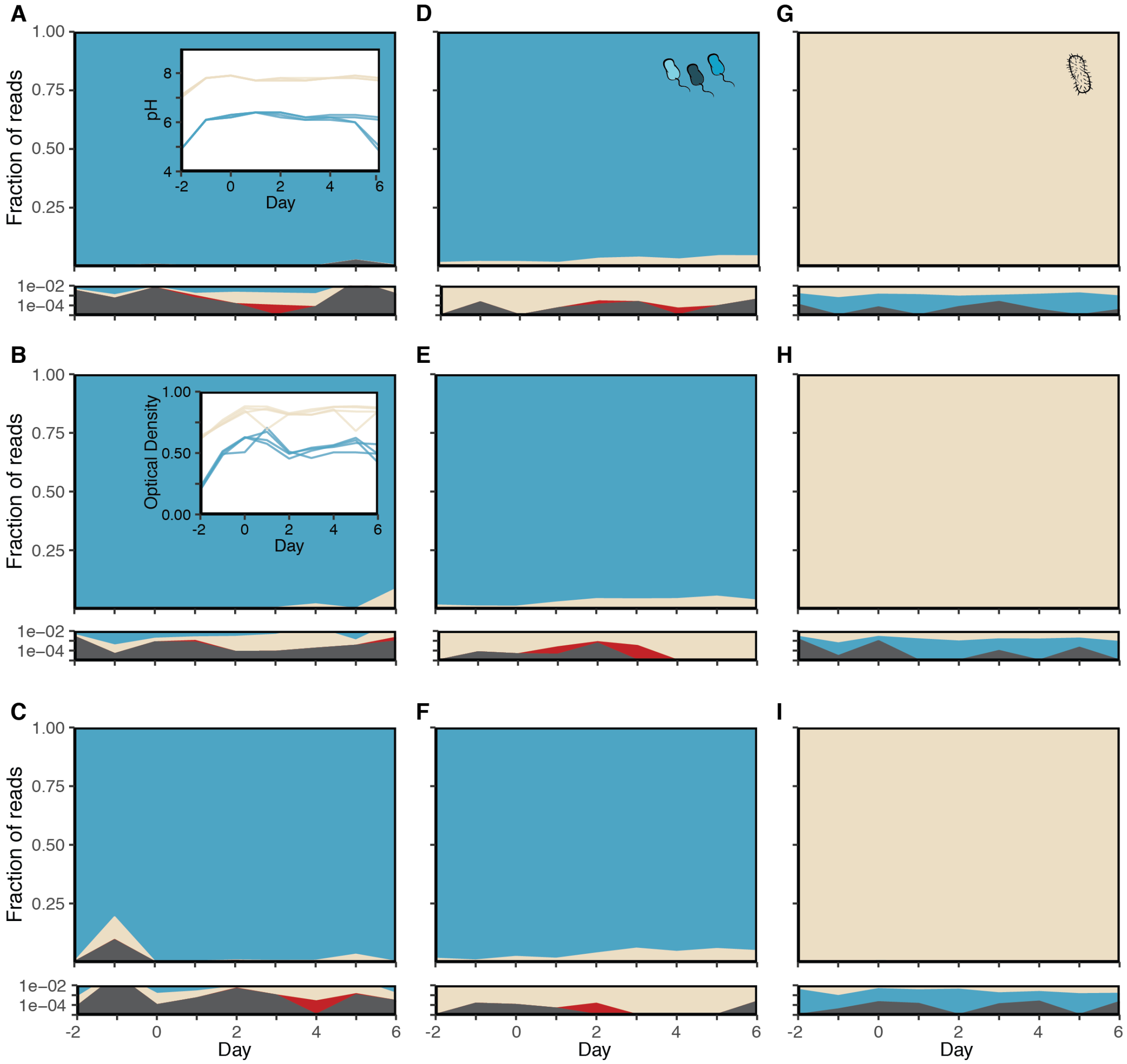


**Fig. S9.**

**Mutual resilience against migration between the *Bacillus* and *Pantoea* soil communities. A-C** Time series for the fraction of 16S reads in 3 technical replicates in which the community dominated by Bacillus (blue) was exposed to migration by Pantoea (cream). Reads from *Pseudomonas* (red) and other (gray) sequence variants were occasionally detected at lower abundances (lower panels, shown in logarithmic scale). We sequenced 3 out of 4 technical replicates, for which pH and optical density measurements are shown in the insets of panels A and B, respectively, along with the same measurements for 4 replicates in which *Pantoea* was exposed to migration by *Bacillus* (lines in cream color in the insets of A and B). The data in A-C correspond to an independent experiment from the one in Fig. 3B. **D-I.** Fraction of 16S read abundances for cultures shown in Fig. 3B. In D-F, the community dominated by *Bacillus* was exposed to migration from *Pantoea*. In G-I, the community dominated by *Pantoea* was exposed to migration from *Bacillus*. We sequenced 3 out of the 4 replicates shown in Fig. 3B for each case. In all the replicate cultures we observed that the two communities were resilient to migration from each other.

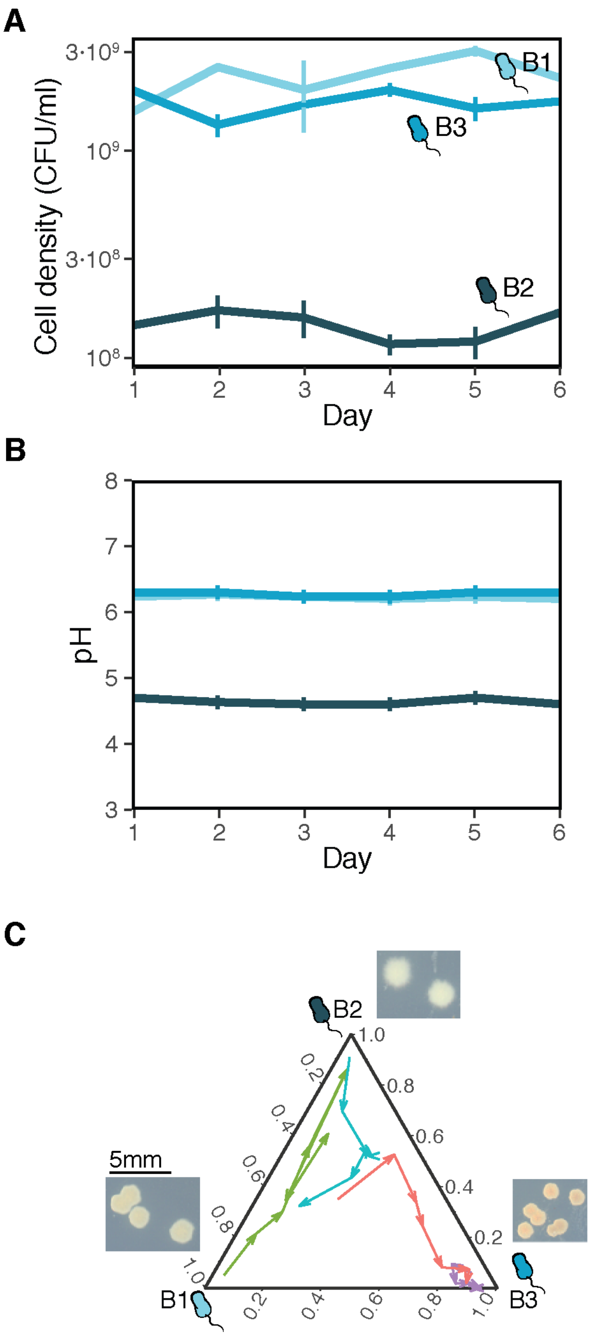


**Fig. S10.**
**Different behavior in laboratory microcosms for 3 genetically similar soil isolates. A.** Time series for the cell density of monocultures from three strains isolated from the soil community (average and standard errors of 3 biological replicates exposed to the 30X daily dilution protocol described in the main text). Strain B2 exhibits a lower carrying capacity than strains B1 and B3. The three strains share the same 16S amplicon sequence, and were found in coexistence in one of the stable states described in Fig. 3. **B.** Time series for the pH (measured at the end of each cycle) of the monocultures in panel A. The strain B2 induces a more acidic pH than the one observed for the other two strains. **C.** Observed trajectories during a competition experiment in which the three isolates were mixed at different initial fractions (different arrow colors) and

exposed to daily dilutions. The three edges of the simplex depict the abundance for the different strains. The observed trajectories indicate that the system can reach two alternative community compositions.

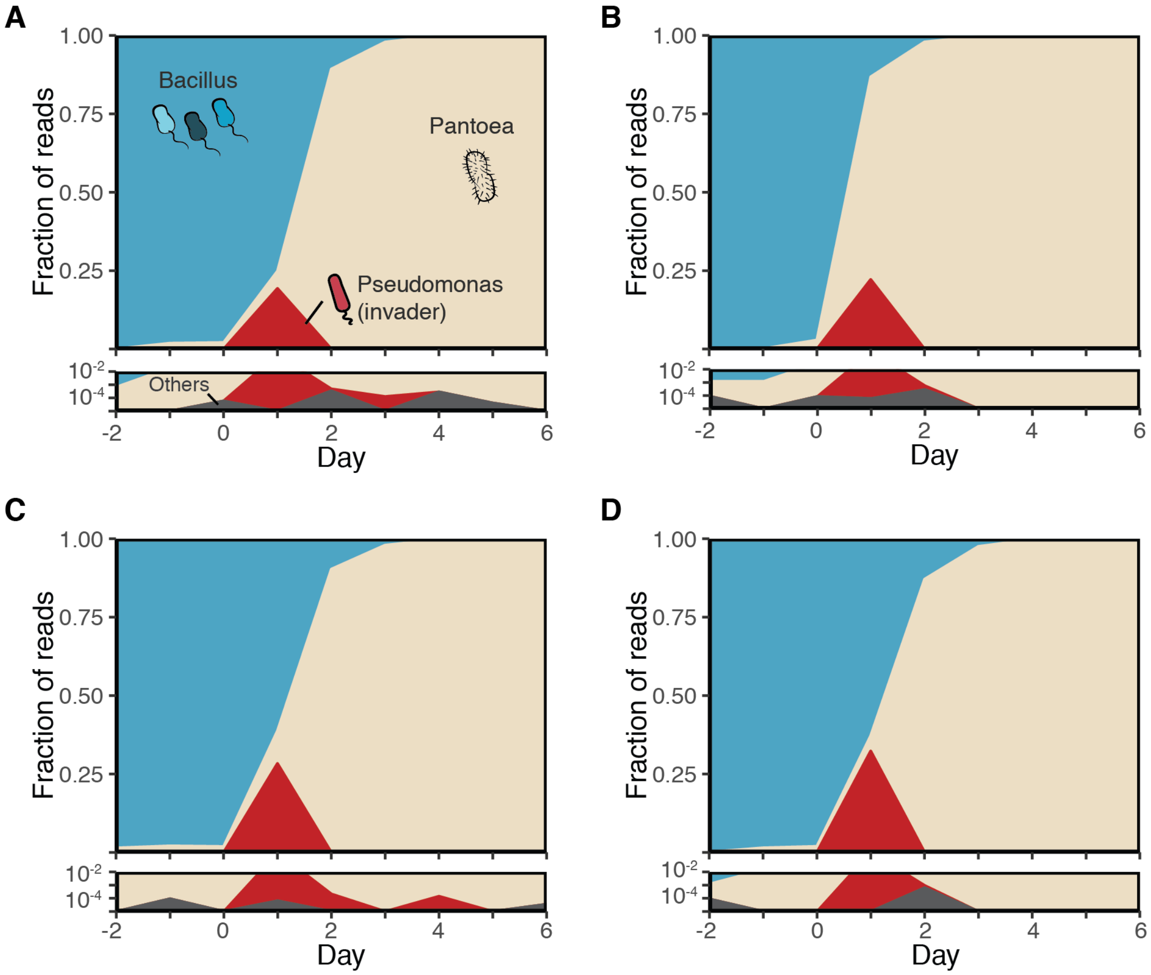


**Fig. S11.**
**Both *Pseudomonas chlororaphis* and *Pseudomonas aurantiaca* can induce transitions from the *Bacillus* to the *Pantoea* stable states. A-B** Time series for the fraction of 16S reads showing the transition from the *Bacillus* stable state to the *Pantoea* stable state following inoculation of *Pc* (two replicates of the experiment shown in Fig. 3C). **C-D** Time series for the fraction of 16S reads showing the transition from the *Bacillus* stable state to the *Pantoea* stable state, in this case following inoculation of *Pa* cells.
